# Microclimatic Effects on Functional Traits of *Arctostaphylos crustacea* ssp. *crustacea* in Alameda County, California, USA

**DOI:** 10.1101/2025.10.03.680375

**Authors:** Leilani G. Hsiao

## Abstract

Anthropogenic climate change and land-use changes threaten the health and survival of plants, particularly by altering the microclimatic conditions of habitats. Plant productivity is highly sensitive to these abiotic conditions that influence their morphological and physiological traits. I studied this relationship in plants of the *Arctostaphylos* genus, commonly known as manzanita, shrubs in chaparral ecosystems across California. I assessed microclimate and examined its effects on leaf morphological traits and plant productivity of *A*. *crustacea* ssp. *crustacea* across two sites in Alameda County, California. At the site of higher sunlight and less soil moisture (VWC), leaves had a greater mass per area (LMA) and were more steeply angled, and photosynthesis was significantly higher (11.14 µmolCO_2_/m^2^s) than leaves at the site of less sunlight and more VWC (7.94 µmolCO_2_/m^2^s). Results from linear mixed models showed that light level was the overall strongest predictor variable for plant traits, with vapor pressure deficit and VWC also contributing to LMA, and leaf temperature and leaf angle distribution also influencing photosynthesis. Overall, no individual microclimatic variable was the sole contributing predictor of a leaf morphological trait or photosynthesis. Rather a combination and interaction of microclimatic conditions influenced plant functional traits, though some conditions had greater influence than others. The functional traits of *A*. *crustacea* ssp. *crustacea* adjusted in response to microclimatic factors, showing intraspecific trait variation (ITV) of this species. ITV is an essential defense for plant resilience that allows for adaptations in the face of rapidly changing climatic conditions.

## INTRODUCTION

Because anthropogenic climate change continues to affect abiotic conditions and biotic dynamics in chaparral ecosystems, there is an increased need to study its effects on plant response. In chaparrals, temperatures are likely to increase, precipitation variability will increase, and droughts will become more frequent or severe or both (Diffenbaugh et al. 2015, Molinari et al. 2018). However, many modeling attempts to predict plant distribution in response to climate change do not incorporate microclimate (Lembrechts et al. 2019, Stark and Fridley 2022, Scherrer and Körner 2011). Microclimate describes the localized climatic conditions within a larger climatic zone, characterized by its distinct physical properties. In chaparral, aspect is one common driver of microclimate. A south-facing slope in the Northern Hemisphere receives more sunlight than north-facing slopes, making them drier and warmer, further affecting plant composition on those slopes (Quinn and Keeley 2006). Such microclimatic variables can potentially moderate climate change effects. For example, de Frenne et al. (2013) found that the shading from a dense forest canopy reduced the abundance of warm–adapted species in the understory. Macroclimate models may be generally applicable to predicting plant response, but a study by Anderegg (2023) found that macroclimate models do not accurately predict plant functional traits due to a lack of microclimatic data.

Another possible contributing factor to poor macroclimatic model prediction of plant functional traits is the lack of intraspecific trait variation (ITV) considered (Anderegg 2023). ITV is the differences in traits within a species across its distribution, reflecting its environmental conditions and allowing individuals to compete with other species (Albert et al. 2011, Clark 2010). High ITV of a species may indicate its ability to survive in high abiotic stress conditions (Umaña et al. 2015). However, ITV is an important yet often not fully studied metric in plant community studies (Westerband et al. 2021, Ahrens et al. 2021). As found in Anderegg (2023), community-level studies fail to fully examine microclimatic effects on plant traits, leading to weak observed relationships between plant traits and environmental conditions.

Plant functional traits, whether morphological, physiological, or phenological, determine how plants interact with their environment and respond to stressors. Various leaf morphological traits are important as they act as a plant’s intermediary to its environment. Stomata on leaves control the uptake of CO_2_ and transpiration of water while leaf angle regulates temperature and adjusts for light interception. These processes may maximize resource acquisition and minimize environmental stress to yield high levels of plant productivity (Fritz et al. 2018, Wright et al. 2004, van Zanten et al. 2010). Leaf morphology changes along environmental gradients (Wang et al. 2022), including that of *Arctostaphylos* (Shaver 1978).

Plants of the *Arctostaphylos* genus, commonly known as manzanita (family *Ericaceae*) are widely recognized for their prolific diversity (100+ overall taxa), including several endemic endangered species, high tolerance to drought, and their ability to grow in the nutrient-poor soils of many California chaparral ecosystems (Kauffmann et al. 2021). One species, widely distributed throughout California, is *Arctostaphylos crustacea* ssp. *crustacea*, commonly known as brittleleaf manzanita. It is a facultative seeder, able to regenerate post-fire both through a seed bank ignited by fire and resprouting from a lignotuber, or burl (Kauffmann et al. 2021). In areas of land-use change where wildfire is suppressed, facultative clonal resprouting of *Arctostaphylos* spp. is common, discouraging possible genetic diversification from a seed bank which would increase the plant’s likelihood of having adaptations to a quickly changing climate (Kauffmann et al. 2021). Further anthropogenic effects, like warming temperatures will prompt increased fuel growth and lower fuel moisture content, creating more favorable conditions for frequent wildfires that will not allow adequate recovery time for Arctostaphylos shrubs to re-establish before the next fire. As the frequency of wildfires increases, woody shrublands will convert to invasive grasslands, destroying ecosystem function and losing biodiversity (Syphard et al. 2019, Jacobsen and Pratt 2018). Although *Arctostaphylos* taxa have co-evolved with chaparral climate and disturbance regimes over the last 15 million years (Parker 2007), rapid climate change and land-use changes threaten *Arctostaphylos* plant health to a new degree.

Current research focuses primarily on the physiological hydraulic traits of *Arctostaphylos* along environmental gradients (e.g. Drake-Schultheis et al. 2020, Vasey et al. 2012). Manzanita are generally considered drought-tolerant, but there is evidence that shallow rooting of *Arctostaphylos*, while the most resistant to cavitation (Pausas et al. 2016), also leaves them the most vulnerable to high-intensity droughts (Pausas et al. 2016, Venturas et al. 2016). Droughts, like the historic 2011-2017 California drought, are becoming increasingly more severe, which in conjunction with other factors, lead to increased stress and mortality of *Arctostaphylos* (Jacobsen and Brandon Pratt 2013, Drake-Schulteis et al. 2020). Water deficit *Arctostaphylos* are more vulnerable to diseases, species competition, abiotic pressures (heat and urbanization), and ultimately mortality. Although studies of hydraulic traits in *Arctostaphylos* species have captured some amount of ITV across the genus, less is known about their intraspecific variation of leaf morphology and productivity. A previous study by Shaver (1978) found that leaf angle and light absorption of *Arctostaphylos* spp. changed along an elevation gradient; however there is a lack of studies examining other climatic conditions or other aspects of leaf morphology.

Plant functional traits, highly sensitive to microclimate, are also indicators of plant response to warming temperatures (Reich et al. 1999, Bruelheide et al. 2018, Auger and Shipley 2013). Therefore it is critical to understand how these plants respond in their functional traits to microclimate and the extent to which they are photosynthetically productive (a measure of gas exchange and carbon assimilation, and proxy for growth), to better predict their response to anthropogenic-driven threats. This study attempts to address this knowledge gap by examining the microclimatic effects on intraspecific variation in plant functional traits in *Arctostaphylos crustacea* ssp. *crustacea*. The study uses two California East Bay Area field sites, a common garden and a natural field site, representing a water availability gradient. Specifically, I ask: **(SQ1)** What are the microclimatic conditions of the study sites?

**(H1)** I hypothesize that the common garden site will have greater water availability than the natural field site but that they will potentially receive similar light levels. Soil temperature will vary based on light, water, and tree canopy, and wind will be similar at both sites.

**(SQ2)** What are the leaf morphological traits and plant photosynthetic productivity of the study species?

**(H2)** I predict that at both sites, leaves will be angled steeply to avoid intense, direct sunlight. If there is a difference in solar radiation between sites, I also hypothesize that the leaf mass per area will be greater at the site of greater solar radiation (Poorter et al. 2009). If a site has greater solar radiation and water availability, I predict that there will be higher productivity in individuals, due to increased resources necessary for photosynthesis.

**(SQ3)** What is the effect of microclimate on the functional traits of the study species?

**(H3)** I predict that solar radiation and water availability will be the most influential microclimatic factors affecting leaf morphology, specifically leaf angle distribution, and plant productivity (Wright et al. 2004).

## METHODS

### Study sites and species

To evaluate the effects of microclimatic conditions, I chose two study sites of varying water availability and unknown, but presumably similar, levels of solar radiation, as both sites are located in the hills of the East Bay in California. The combination of seasonal precipitation, elevation, and soil conditions at each site determined its water availability, allowing me to characterize them as mesic, with sufficient water availability, or xeric, with insufficient water. The mesic site is located within a common garden, the University of California Botanical Gardens (UCBG) in Berkeley, California (37.875°N, 122.238°W). The site regularly receives care from garden staff and is watered during the dry season (June-September) every 3-4 weeks for 1-3 hours per watering event. Huckleberry Botanic Regional Preserve (HRP) is a xeric field site in Oakland, CA (37.843°N, 122.194°W). A natural area within the East Bay Regional Park District (EBRPD), HRP does not receive additional water aside from natural fog and rainfall.

The University of California Berkeley Botanical Gardens is a 34-acre plot of land, housing several “ecoregions” of plants from regions around the world, including California. UCBG has 257 recorded individuals within the *Arctostaphylos* genus in different sections of its Californian collection. Study individuals of *A. crustacea* ssp. *crustacea* were located in Bed 26, Accession 82.1644, originally obtained from an unknown California location; Bed 27, Accession 83.0009 from Monterey, CA; and Bed 27, Accession 82.1601 from Contra Costa, CA.

In HRP, soil conditions are characterized by bands of interbedded chert and shale, gravelly soil, low nutrient values, and poor water holding capacity in shallow soils (East Bay Regional Park District). Study organisms were located at the “Manzanita Barren,” as indicated by EBRPD’s map, accessible by the Interpretive Loop trail. The endangered *A. pallida* and *A. crustacea* ssp. *crustacea* dominate this preserve. I sampled three individuals of the non-endangered but endemic *A. crustacea* ssp. *crustacea* from the Manzanita Barren. To mitigate the likelihood of the samples being clones, I chose three individuals geographically separate from each other, with *A*. *pallida* individuals interspersed.

I chose the study species, *A. crustacea* ssp. *crustacea,* based on its extensive distribution in California, availability in the study sites, and accessibility on the trails. Species were clearly labeled at UCBG and easily distinguishable at HRP. To further confirm identification, I identified species using a field guide (Kauffmann et al. 2021), as well as a hand lens to examine the plant.

### Microclimate

Sampling was conducted from early February to late March 2025 in the mornings, from 09:00 to 12:00. I chose sunny days with no predicted rain forecast as a time of likely higher plant productivity. Due to instrument availability and weather conditions, UCBG data was collected in the beginning to mid-February and HRP data was collected at the end of March, influencing the microclimates. These microclimatic measurements were taken simultaneously with leaf morphology and productivity measurements.

I conducted various measurements to assess the microclimate of each site (Table 1). First I calculated vapor pressure deficit (VPD) as defined by Abtew and Melesse (2012), using measurements of temperature and relative humidity taken by an EL-USB-2+ data logger (Lascar Electronics, Essex, England) and the local atmospheric pressure as measured by local weather stations (Visual Crossing Weather, Weather Underground). Data loggers were placed on branches of the study individuals to record exact temperature and relative humidity. Using a soil moisture sensor (Decagon Devices, ECH20 Model EC-5, Meter Group Inc. Pullman, WA, USA), I measured soil moisture as volumetric water content (VWC). To measure solar radiance given a leaf’s specific angle, I used a light sensor on a LI-6800 Portable Photosynthesis System (LI-COR, Inc. Lincoln, NE, USA). Additionally I used a Kestrel 1000 Wind Meter to measure wind speed (Kestrel Instruments, Nielsen-Kellerman Co., Boothwyn, PA, USA).

**Table 1.** Measurements used in the study, their abbreviations, units, and purpose.

| Abbreviation | Units | Measurement | Purpose |
| --- | --- | --- | --- |
| VPD | kPa | Vapor pressure deficit | Microclimate |
|  | m | Elevation | Microclimate |
|  | m/s | Wind speed | Microclimate |
| VWC | % | Volumetric water content of soil | Microclimate |
| T<sub>soil</sub> | °C | Temperature of the soil surrounding the plant | Microclimate |
| Q | μmol/m<sup>2</sup>/s | Amount of solar radiation the leaf is receiving as photosynthetic photon flux density | Microclimate |
|  |  | Aspect, the compass direction a slope faces: north, east, south, and west | Microclimate |
| Leaf length | mm | Length from base to tip of leaf | Leaf morphology |
| Leaf width | mm | Width at the widest point of leaf | Leaf morphology |
| Leaf height | cm | Height at which the leaf is located on the plant relative to total plant height | Leaf morphology |
| Plant height | cm | Total height of plant | Leaf morphology |
| Plant width | cm | Width of plant at the widest part | Leaf morphology |
| T<sub>leaf</sub> | °C | Temperature of the leaf | Leaf morphology |
| LAD | ° | Leaf angle distribution of the whole plant | Leaf morphology |
| LMA | mg/cm<sup>2</sup> | Leaf mass per area, a measure of biomass invested per unit leaf area | Leaf morphology |
| A | μmolCO<sub>2</sub>/m<sup>2</sup>s | Rate of carbon assimilation of leaf | Plant productivity |

### Leaf morphology

To characterize leaf morphology of each species, I conducted physical measurements of 3 leaves per plant per site (Table 1). Using a digital Mitutoyo caliper with an accuracy of ±0.02 mm (Model 500-196-30, Mitutoyo Corporation, Kanagawa, Japan), I measured leaf length and width. I documented leaf temperature with a FLIR E54 thermal camera (Teledyne Flir, FLIR E54 24°, Teledyne FLIR LLC, Wilsonville, OR, USA). Leaves were chosen based on their maturity and health: green, non-blighted leaves on a second-year stem to optimize plant productivity.

To determine leaf mass per area (LMA) using 3 leaves from each plant (after measuring their productivity as described below), I photographed them against a 5×5 cm square, calculated leaf area in cm^2^ using the app Leafscan (Carlos Anderson 2017), and oven-dried the leaves at 60°C for 48 hours to weigh their dry mass in grams. At HRP, the brittleleaf leaves were true to their name, so I measured them after quantifying their rates of productivity.

To obtain leaf angle distribution (LAD), I took photographs using an iPhone 13 of the plant canopy with a leveled tripod at surrounding angles (north, east, south, west). If the study individuals were blocked by other plants, I took photographs from ordinal directions. For each direction, I took at least 4 photographs. As per Ryu et al. (2010), I chose leaves oriented parallel to the viewing direction of the camera and used ImageJ to obtain leaf angles. I measured 20 leaf angles from each direction for each plant, totaling 80 leaf angles per plant. I used package ‘*LAD*’ by Francesco Chianucci (2022) to plot LAD and obtain mean leaf inclination angle and distribution type.

I also measured plant height and width using a yardstick to estimate plant size.

### Plant productivity

To determine plant productivity, I measured photosynthesis as carbon assimilation (A) with a LI-6800 Portable Photosynthesis System between 09:00 and 12:00 under sunny, non-cloudy conditions (Table 1). I used these same leaves for leaf morphology measurements. I took at least 3 readings per leaf, using 3 leaves distributed throughout the canopy of each plant. I measured 3 plants at each site.

### Data Analysis

To determine the significance of microclimatic variables on leaf morphological traits and plant productivity, I used various statistical analyses methods with R Statistical Software (v4.5.0; R Core Team 2025). I used packages ggplot2 (Wickham 2009), dplyr (Wickham et al. 2023), lme4 (Bates et al. 2015), DHARMa (Hartig 2024), and flexplot (Fife 2022).

I used t-tests, analysis of variance (ANOVA), and post hoc Tukey tests to assess statistical significance in microclimate, leaf morphology, and plant productivity. To answer SQ1 and SQ2, I compared mean VWC, Q, VPD, T_soil_, wind speed, leaf mass per area (LMA), leaf angle distribution (LAD), and photosynthesis across the study sites. I also ran linear mixed models (LMMs) to test if microclimatic variables caused any variation in leaf morphological traits and plant productivity to address SQ3 (Table 2). Fixed effects were microclimatic metrics, some interactive (indicated by * between metrics) while others were non-interactive (indicated with + between metrics). The location and plants were used as random effects. I ran LMMs with both single and multiple predictor variables. The exact formulas were chosen based on results from relationships between microclimate and leaf morphology or plant productivity.

**Table 2.** Linear mixed model formulas used to assess microclimatic effects on plant functional traits. Outcome variables were LMA, LAD, and A; predictor variables were individual microclimatic variables: VWC, Q, VPD, T_soil_, and wind speed; random effect was location (HRP or UCBG) or individual plants.

|  |  |
| --- | --- |
| **Single microclimatic variable** | LMA~VWC+(1\|location)<br>LMA~Q+(1\|location)<br>LMA~VPD+(1\|location)<br>LMA~$T_{\text{soil}}$+(1\|location)<br>LMA~wind+(1\|location)<br>LAD~VWC+(1\|location)<br>LAD~Q+(1\|location)<br>LAD~VPD+(1\|location)<br>LAD~$T_{\text{soil}}$+(1\|location)<br>LAD~wind+(1\|location)<br>A~VWC+(1\|location)<br>A~Q+(1\|location)<br>A~VPD+(1\|location)<br>A~$T_{\text{soil}}$+(1\|location)<br>A~wind+(1\|location) |
| **Multiple microclimatic variables** | Leaf length~scale(Q)+VPD+plant height+(1\|plant)<br>LMA~scale(Q)+VWC+VPD+(1\|plant)<br>A~scale(Q)+VWC+VPD+plant height+(1\|plant)<br>A~scale(Q)*LAD+(1\|plant)<br>A~scale(Q)*$T_{\text{leaf}}$+LAD+(1\|plant) |

## RESULTS

### Microclimate

UCBG and HRP differed in all microclimatic variables: VWC, Q, VPD, T_soil_, and wind speed. Between sites, all mean microclimatic measurements were statistically significantly different (p-value<0.05) (Table 3, Table 4). Overall, UCBG experienced lower mean solar radiation (Q) (896.93 µmol/m^2^s), greater mean wind speed (0.85 m/s), and cooler mean soil temperatures (21.17°C) than HRP (Figure 2, Table 3). A potential influence on these cool climatic conditions was the presence of several pine and oak trees above the study individuals, shading the understory. UCBG also had greater mean VWC (25.29%) and a lower mean VPD (1.28 kPa) compared to HRP, suggesting relatively high water availability in the soil and air (Table 3). Plants at HRP were exposed to greater mean solar radiation (1734.43 µmol/m^2^s) and a slower mean wind speed (0.39 m/s) than at UCBG (Figure 2, Table 4). HRP experienced a low mean VWC (6.6%) and a large range of VPD (mean 2.06 kPa), and a range of soil temperatures (mean 25.88°C) (Figure 2, Table 4). The coarse-textured soil and presence of fog likely influenced the water availability properties of HRP (EBRPD).

**Table 3.** UCBG mean microclimatic variables and standard deviations. n=45 observations. *** denotes a p-value from [0, 0.001], ** denotes a p-value from (0.001, 0.01), and * denotes a p-value from (0.01, 0.05] from a t-test comparison of variables between sites (Table 3 and Table 4).

| Microclimatic Variable | Units | Mean value | SD |
| --- | --- | --- | --- |
| Q | $\mu\text{mol}/\text{m}^2/\text{s}$ | 896.93*** | 45 |
| VPD | kPa | 1.28*** | 0.369 |
| VWC | % | 25.29*** | 7.91 |
| Soil temperature | $^{\circ}\text{C}$ | 21.17** | 2.82 |
| Wind speed | m/s | 0.85*** | 0.297 |

**Table 4.** HRP mean microclimatic variables and standard deviations. n=69 observations.

| Microclimatic Variable | Units | Mean value | SD |
| --- | --- | --- | --- |
| Q | $\mu\text{mol}/\text{m}^2/\text{s}$ | 1734.43*** | 250 |
| VPD | kPa | 2.06*** | 1.16 |
| VWC | % | 6.60*** | 1.58 |
| Soil temperature | $^{\circ}\text{C}$ | 25.88** | 9.32 |
| Wind speed | m/s | 0.39*** | 0.372 |

**Figure 1.**
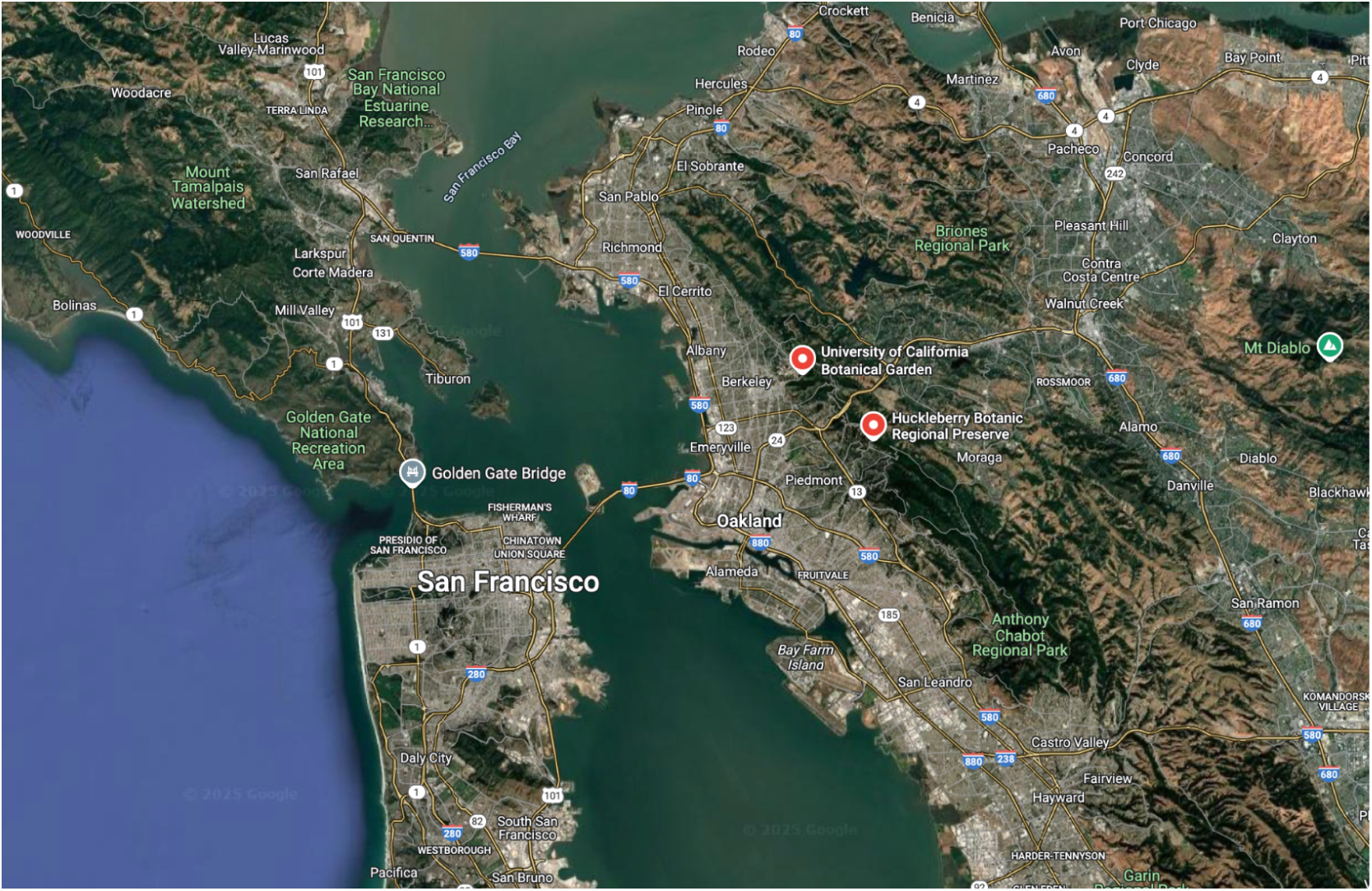
Map of the two study sites, obtained from Google Maps. Including (1) UC Berkeley Botanical Gardens (2) Huckleberry Botanic Regional Preserve.

**Figure 2.**
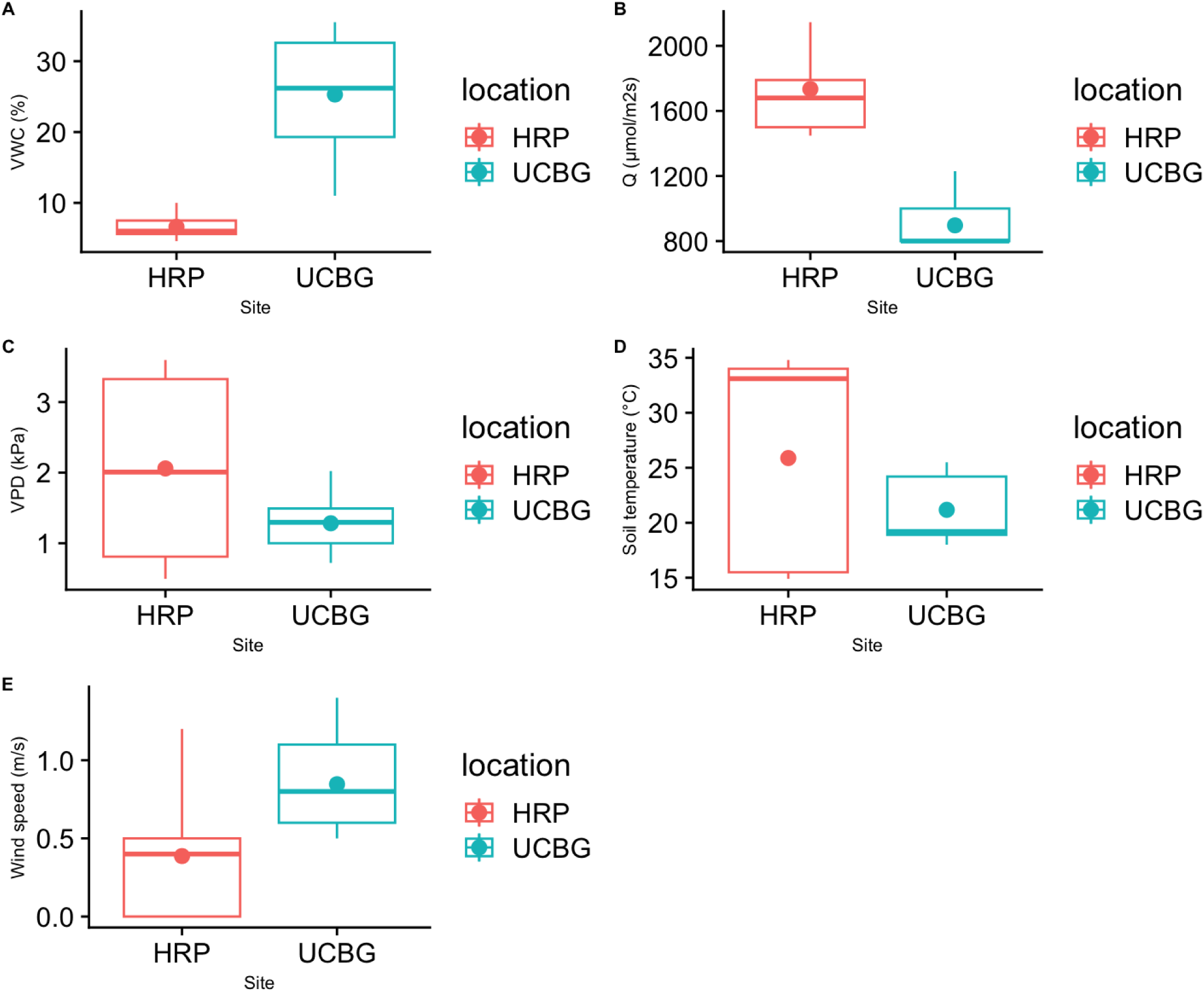
Microclimatic variables between sites. The means of all variables were statistically significantly different between sites. Mean values and standard deviations are found in Table 3. HRP showed a greater range of microclimatic conditions, specifically VPD and soil temperature, than UCBG.

### Leaf morphology

Using a t-test, I found that mean leaf length, mean leaf width, mean LMA, mean LAD, and mean T_leaf_ differed significantly between sites (Table 5, Table 6).

**Table 5.**
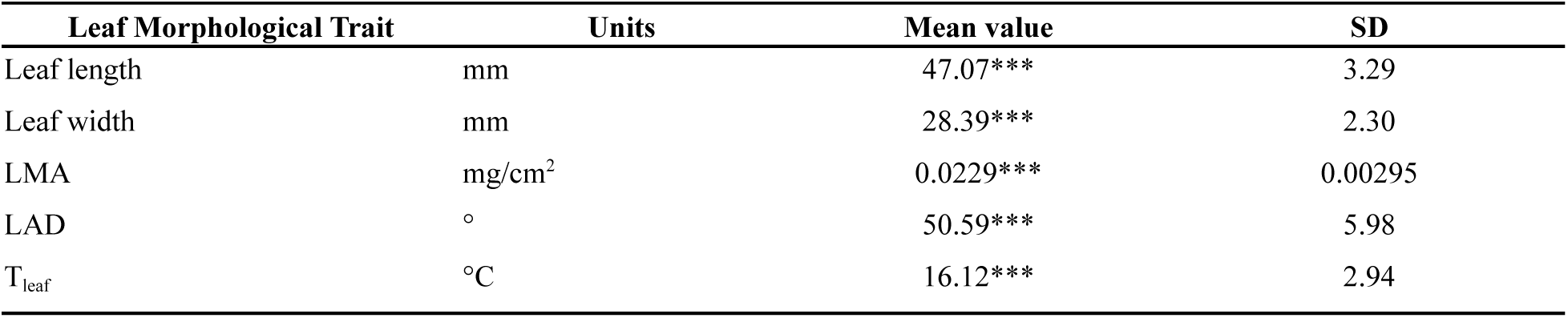
Mean leaf morphological traits at UCBG. n=9 leaves except for LAD, n=240. A t-test was used to compare traits between sites (Table 5 and Table 6).

| Leaf Morphological Trait | Units | Mean value | SD |
| --- | --- | --- | --- |
| Leaf length | mm | 47.07*** | 3.29 |
| Leaf width | mm | 28.39*** | 2.30 |
| LMA | $\text{mg}/\text{cm}^2$ | 0.0229*** | 0.00295 |
| LAD | $^{\circ}$ | 50.59*** | 5.98 |
| $T_{\text{leaf}}$ | $^{\circ}\text{C}$ | 16.12*** | 2.94 |

**Table 6.** Mean leaf morphological traits at HRP. n=9 leaves except for LAD, n=240. A t-test was used to compare traits between sites (Table 5 and Table 6).

| Leaf Morphological Trait | Units | Mean value | SD |
| --- | --- | --- | --- |
| Leaf length | Mm | 39.52*** | 3.71 |
| Leaf width | Mm | 26.88*** | 1.31 |
| LMA | $\text{mg}/\text{cm}^2$ | 0.0295*** | 0.00177 |
| LAD | $^{\circ}$ | 56.46*** | 5.03 |
| $T_{\text{leaf}}$ | $^{\circ}\text{C}$ | 22.28*** | 1.64 |

Mean LMA was significantly greater at HRP (0.0295 mg/cm^2^) than at UCBG (0.0229 mg/cm^2^). Longer (47.07 mm) and wider (28.39 mm) leaves at UCBG than HRP (39.52 mm, 26.88 mm) increased leaf area, subsequently leading to a smaller LMA at UCBG (Table 5, Table 6). Individuals at both sites had an overall spherical leaf angle distribution (Figure 3, Figure 4), with leaves equally likely to be angled horizontally or vertically. Leaf temperature also differed significantly between sites, with HRP having a greater mean leaf temperature (22.28°C) than UCBG (16.12°C) (Table 5, Table 6, Figure 7).

**Figure 3.**
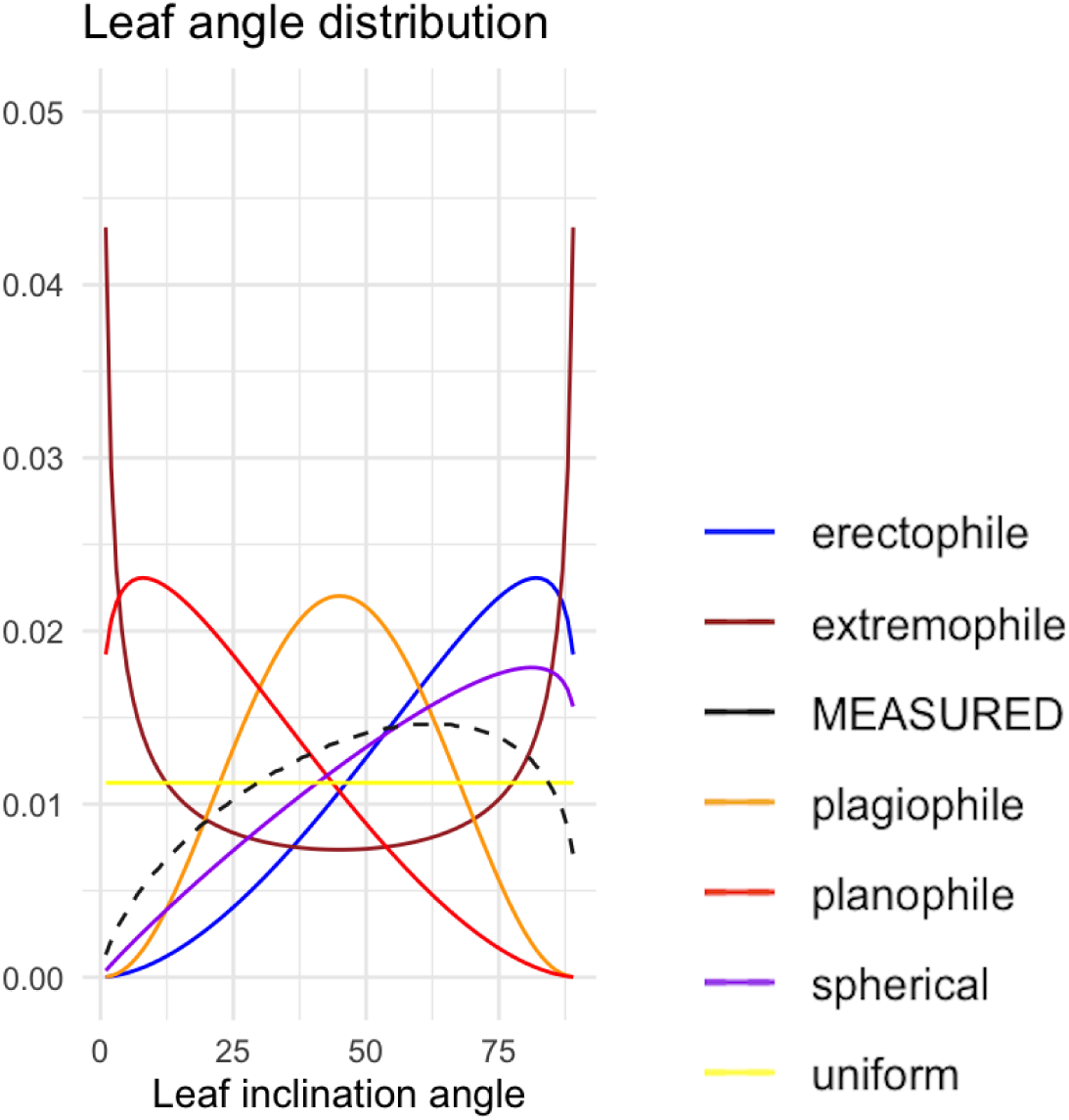
Leaf angle distribution (LAD) at UCBG. The graph shows the distribution of leaf angles for theoretical distributions. The dotted black line represents the LAD for the measured data at UCBG; in this case, fitting most closely to the spherical distribution (n=240 observations).

**Figure 4.**
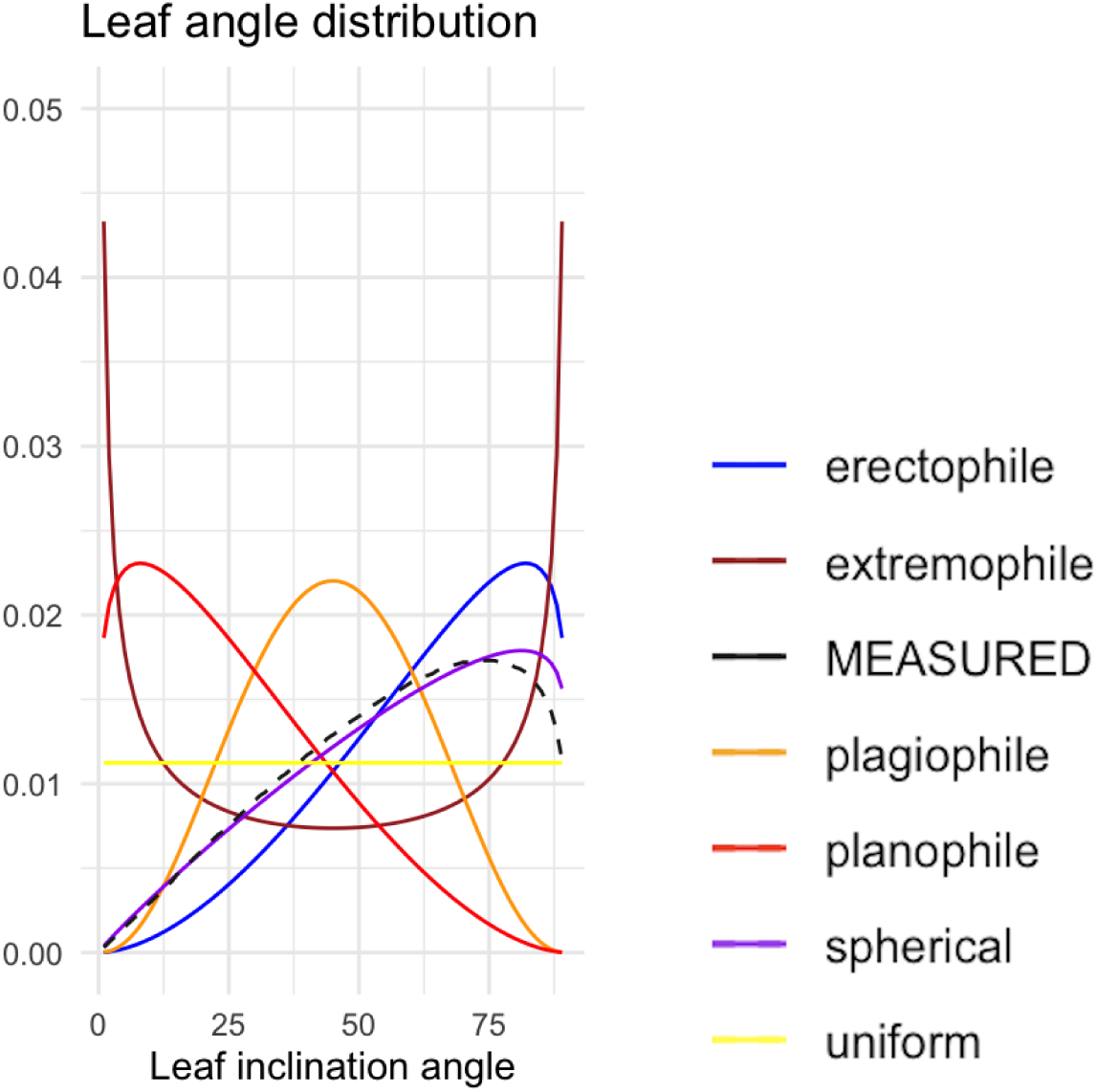
Leaf angle distribution (LAD) at HRP. The black, dotted line of the measured LAD at HRP fits closely with the spherical distribution ( n=240 observations).

**Figure 5.**
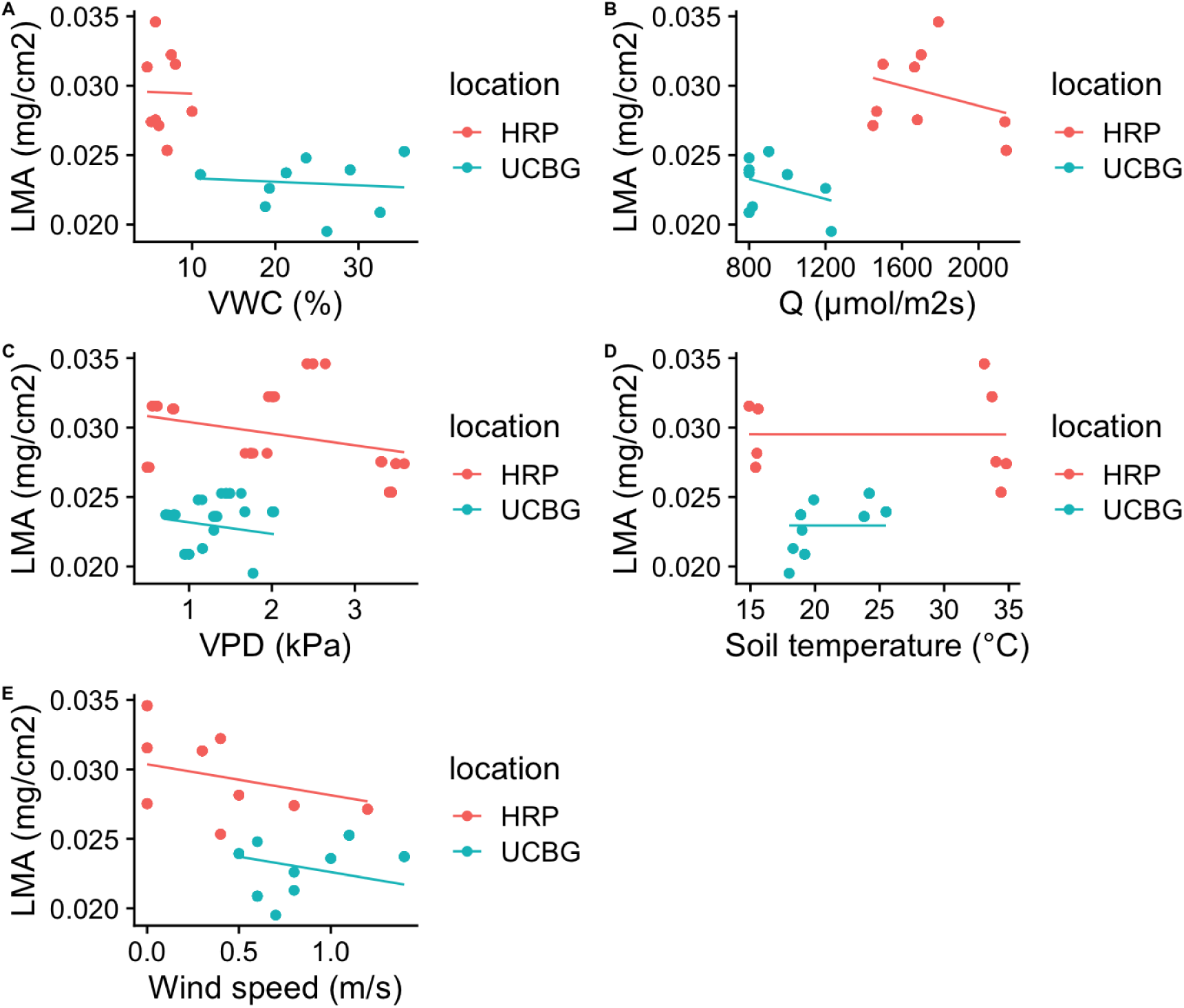
Microclimatic variables and LMA across sites. LMA means were statistically significantly different between sites. Mean values and standard deviations are found in Table 5 and Table 6.

When LMA was correlated with an individual microclimatic variable, I found that there were mostly very weak, if any, correlations present (Table 7). Of the microclimatic variables, there was a significant p-value (<0.05) for Q, VPD, and wind speed acting on LMA (Table 7). However the R^2^m values were low and R^2^c-R^2^m values were high; for example, the R^2^m of Q was about 0.05 and R^2^c-R^2^m was about 0.84 (Table 7), meaning only 5% of the variance in LMA is explained by the fixed effect, Q, while 84% of the variance in data is due to the random effect, location. I also compared mean LMA across different aspects (north, south, west) at both sites (Figure 6). A one-way ANOVA yielded a non-significant difference in mean LMA across aspects at each site (p-value=0.841). However, a post hoc Tukey test yielded a p-value of 0 for south-and north-facing slopes and for west-and north-facing slopes and a p-value of 0.0182 for west- and south-facing slopes at HRP. At UCBG, a Tukey test resulted in a nonsignificant p-value of 0.1677 for west- and south-facing slopes.

**Figure 6.**
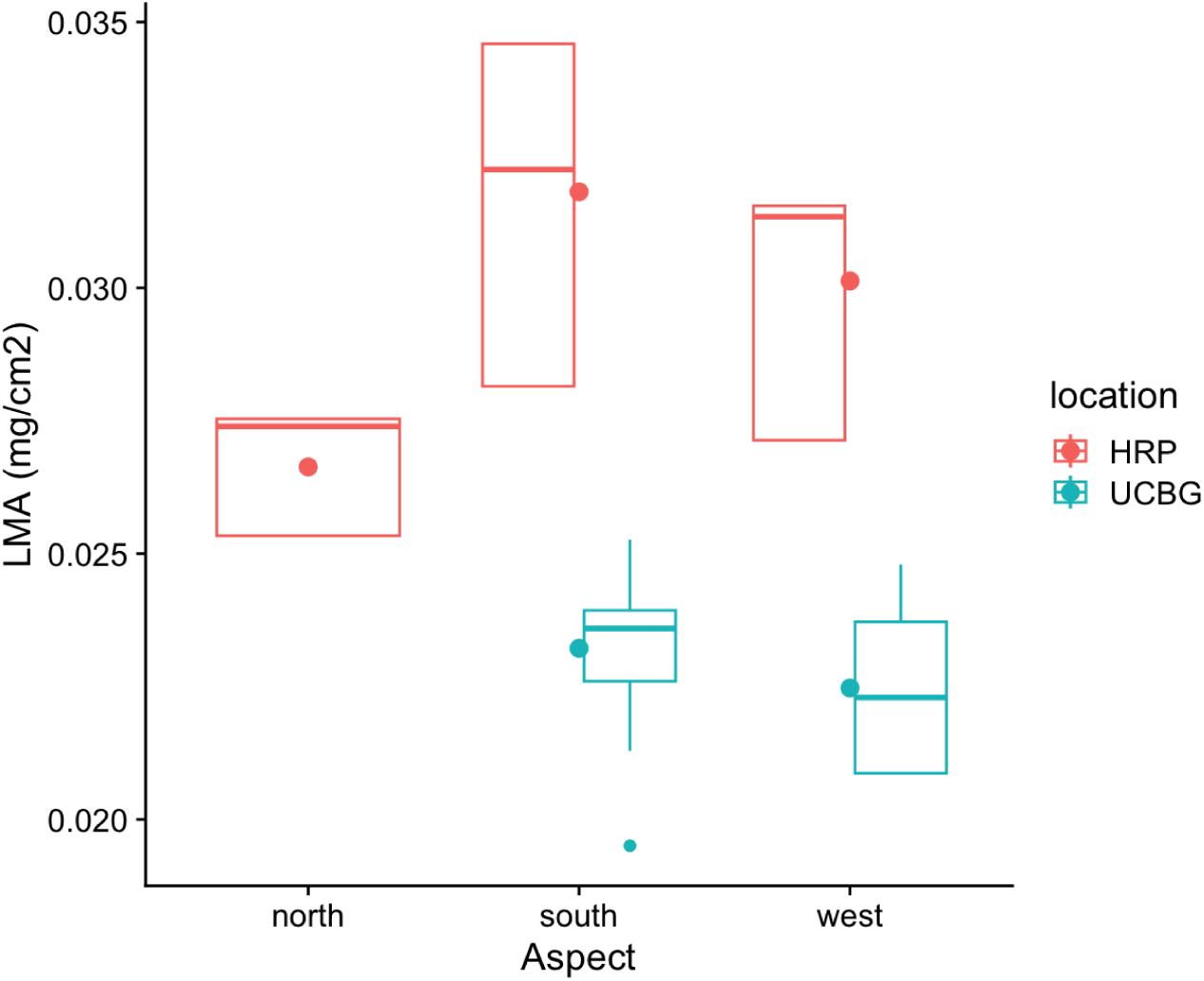
Aspect and LMA across sites. LMA was significantly different between sites. At HRP, LMA were significantly different between all aspects. At UCBG, LMA was not significantly different between aspects.

A similar trend occurred with the correlations between microclimatic variables and mean LAD. Between sites, there were no significant differences in the weak response LAD displayed to microclimatic variables (Figure 8). Only Q and VPD showed a statistically significant p-value, yet all linear mixed models for mean LAD yielded relatively high variances explained by random effects, ranging from about 30% for VWC to about 65% for Q (Table 7). Using a one-way ANOVA, I compared mean LAD across different aspects (north, south, west) at both sites. Results showed a significant difference in mean LADs across aspects at each site (p-value=1.4e-12). After a post hoc Tukey test, I found that the mean LAD at each aspect was significantly different (p-value=0) from another at each site. At HRP, mean LAD at south- and west-facing slopes were significantly higher than mean LAD at north-facing slopes (Figure 9). At UCBG where plants were only present on a south- or west-facing slope, mean LAD at south-facing slopes was significantly higher than mean LAD at a west-facing slope (Figure 9).

**Table 7.** Linear mixed model formulas, p-value, marginal R^2^, conditional R^2^, and difference between R^2^s. Marginal R^2^ values (R^2^m) signify the variance in outcome variable data explained by the fixed effects, conditional R^2^ values (R^2^c) signify the variance in outcome data explained by the entire model (fixed and random effects), and the difference (R^2^c-R^2^m) is the variance in outcome data explained by random effects.

| Model formulas used in *lme4* notation | p-value | $R^2_m$ | $R^2_c$ | $R^2_c - R^2_m$ |
| --- | --- | --- | --- | --- |
| LMA~VWC+(1\|location) | 0.5839 | 0.0028 | 0.7438 | 0.741 |
| LMA~Q+(1\|location) | 0.0007*** | 0.0508 | 0.8930 | 0.8422 |
| LMA~VPD+(1\|location) | 0.0006519*** | 0.0217 | 0.819 | 0.7974 |
| LMA~T<sub>soil</sub>+(1\|location) | 0.0008715*** | 0.0377 | 0.7312 | 0.6936 |
| LMA~wind+(1\|location) | 0.9778 | 1.7540e-06 | 0.7675 | 7.67e-01 |
| LAD~VWC+(1\|location) | 0.5723 | 0.0073 | 0.3074 | 0.3001 |
| LAD~Q+(1\|location) | 0.002191** | 0.0965 | 0.7455 | 0.6490 |
| LAD~VPD+(1\|location) | 0.01715* | 0.0312 | 0.4673 | 0.4362 |
| LAD~T<sub>soil</sub>+(1\|location) | 0.388 | 0.0063 | 0.3199 | 0.3135 |
| LAD~wind+(1\|location) | 0.335 | 0.0055 | 0.3905 | 0.3851 |
| A~VWC+(1\|location) | 0.2672 | 0.0154 | 0.6580 | 0.6426 |
| A~Q+(1\|location) | $<2.2e-16$*** | 0.6796 | 0.6796 | 0 |
| A~VPD+(1\|location) | 0.000114*** | 0.0451 | 0.7065 | 0.6614 |
| A~T<sub>soil</sub>+(1\|location) | 0.3632 | 0.0031 | 0.7038 | 0.7007 |
| A~wind+(1\|location) | 0.03395* | 0.0126 | 0.7109 | 0.6983 |

### Plant productivity

Overall, mean carbon assimilation was significantly higher at HRP (11.14 µmolCO_2_/m^2^/s) than at UCBG (7.94 µmolCO_2_/m^2^/s) (Table 8). The predictor effects of Q, VPD, and soil temperature were significant with A, however variance explained by random effects was high for VPD and soil temperature (about 66% and 69%, respectively) (Table 7). For Q, the variance from random effects was 0%, and the variance in A explained by Q was about 67% (Table 7).

**Table 8.**
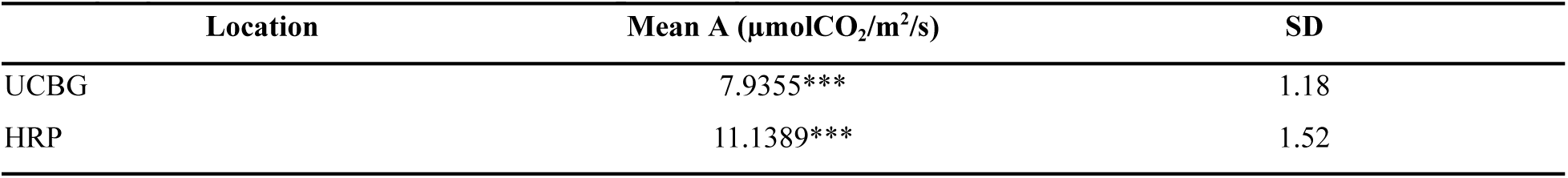
Average Carbon Assimilation (A) Values. A minimum of 3 readings were taken for each leaf, and 3 leaves per plant were measured. n=114 readings. Both p-values from a t-test <2.2e-16.

| Location | Mean A ($\mu\text{molCO}_2/\text{m}^2/\text{s}$) | SD |
| --- | --- | --- |
| UCBG | 7.9355*** | 1.18 |
| HRP | 11.1389*** | 1.52 |

Carbon assimilation also varied with aspect (Figure 11). A one-way ANOVA test showed A differing significantly across aspects at each site (p=5.29e-9). A post hoc Tukey test showed that at HRP, A differed significantly between south- and north-facing slopes (p=0.0003), significantly between west- and north-facing slopes (p=0.0072), and non-significantly between south- and west-facing slopes (p=0.5511). At UCBG, A differed significantly between the south- and west-facing slopes (p=0.0084).

### Microclimatic predictors and interactions

Because no single microclimatic variable had a strong predicting effect on leaf morphology or plant productivity traits, I ran LMMs with multiple predictor variables. Although there were many possible combinations of predictor variables, I chose the model formulas based on the results from relationships between microclimate and leaf morphology or plant productivity.

For leaf length, I found the strongest predictor variables were Q, VPD, and plant height (Table 9). The variance in leaf length data explained by fixed effects was about 71%, while the variance explained by random effects was about 2% (Table 9). For LMA, I found the strongest predictor variables were VWC and VPD, with about 40% of the variance in LMA data explained by fixed effects and 54% of the variance explained by random effects (Table 9). For A, Q and T_leaf_ were the strongest predictor variables, and about 79% of the variance in A data was explained by the fixed effects, while only about 3% were explained by random effects (Table 9).

**Table 9.** Formulas used in linear mixed models, with p-values for each predictor variable, marginal R^2^, conditional R^2^, and difference between R^2^s. The predictor variables were chosen based on previous correlations and the best fit of data.

| Model formulas used in *lme4* notation | p-value | $R^2_m$ | $R^2_c$ | $R^2_c - R^2_m$ |
| --- | --- | --- | --- | --- |
| leaf length~scale(Q)+VPD+plant height+(1\|plant) | Q: < 2.2e-16 ***<br>VPD: 5.268e-06 ***<br>Height: 8.883e-05 *** | 0.7133 | 0.7410 | 0.0277 |
| LMA~scale(Q)+VWC+VPD+(1\|plant) | Q: 0.72468<br>VWC: 0.00218 **<br>VPD: 8.767e-06 *** | 0.4012 | 0.9426 | 0.5415 |
| A~scale(Q)+VWC+VPD+plant height+(1\|plant) | Q: 0.000295***<br>VWC: 0.312498<br>VPD: 0.379945<br>Height: 0.021933 * | 0.6693 | 0.7280 | 0.0587 |
| A~scale(Q)*LAD+(1\|plant) | Q: 4.833e-09 ***<br>LAD: 0.9608<br>Q:LAD: 0.7931 | 0.5569 | 0.6884 | 0.1315 |
| A~scale(Q)*T<sub>leaf</sub>+LAD+(1\|plant) | Q:< 2.2e-16 ***<br>T<sub>leaf</sub>: 0.0011662 **<br>LAD: 0.0008109 ***<br>Q:T<sub>leaf</sub>: < 2.2e-16 *** | 0.7985 | 0.8357 | 0.0371 |

## DISCUSSION

In this study, I examined how leaf morphological and physiological traits responded to microclimatic factors. By assessing microclimate (SQ1), characterizing leaf morphology, and measuring plant productivity (SQ2), I found there was significant variation in the plant traits in response to certain microclimatic variables (SQ3)-notably solar radiation, VWC, and VPD. Overall, individuals presented a spherical LAD, with individuals at UCBG having more horizontally angled leaves than those at HRP. LMA differed slightly between sites. Carbon assimilation showed slight positive correlations with Q and VPD, while there were significant differences in A across a water availability gradient. In agreement with my hypotheses, variation in these morphological and physiological traits differed depending on environmental factors, suggesting that microclimate plays a strong role in driving intraspecific trait variation.

### Functional traits response to microclimate

In agreement with H2 and H3, all leaf morphological traits differed significantly between sites and responded differently to microclimatic variables.

Results showed that higher LMA values were associated with south-facing slopes and lower LMA values with north-facing slopes, suggesting that individuals with thicker leaves favor south-facing slopes (Figure 6). This trend is consistent with previous studies that found that leaves on south-facing slopes, which receive more sunlight, have a high LMA, with a greater leaf thickness and smaller area (Ackerly et al. 2002, Wright et al. 2004). Conversely, north-facing slopes are correlated with a low LMA, with leaves having a larger area to compensate for less sunlight exposure (Ackerly et al. 2002). Lower LMA at UCBG, the site of less solar radiation, and significantly lower LMA on north-facing slopes than south- and west-facing slopes at both sites suggest that light is a limiting resource and that plants will increase leaf area to intercept more sunlight. However, the LADs and leaf temperature suggest that there was also a limiting threshold of radiation. Plants at HRP had leaves angled more steeply and lower photosynthesis on south-facing slopes (Figure 9, Figure 11), while leaf temperature was high on north- and south-facing slopes (Figure 7). Steeper leaf angles are thought to decrease solar exposure when the sun is at high angles, decreasing the chance of photoinhibition and stress on a plant (Falster and Westoby 2003, van Zanten et al. 2010), as well as minimizing transpiration (Cowan 1982). Higher photosynthesis on north-facing slopes and a more horizontal mean leaf angle at HRP (Figure 9, Figure 11) potentially reflect a strategy by the plants to angle leaves horizontally in areas of less sun exposure and to angle leaves vertically on south-facing slopes to avoid the intense sunlight, transpiration, and potential overheating.

**Figure 7.**
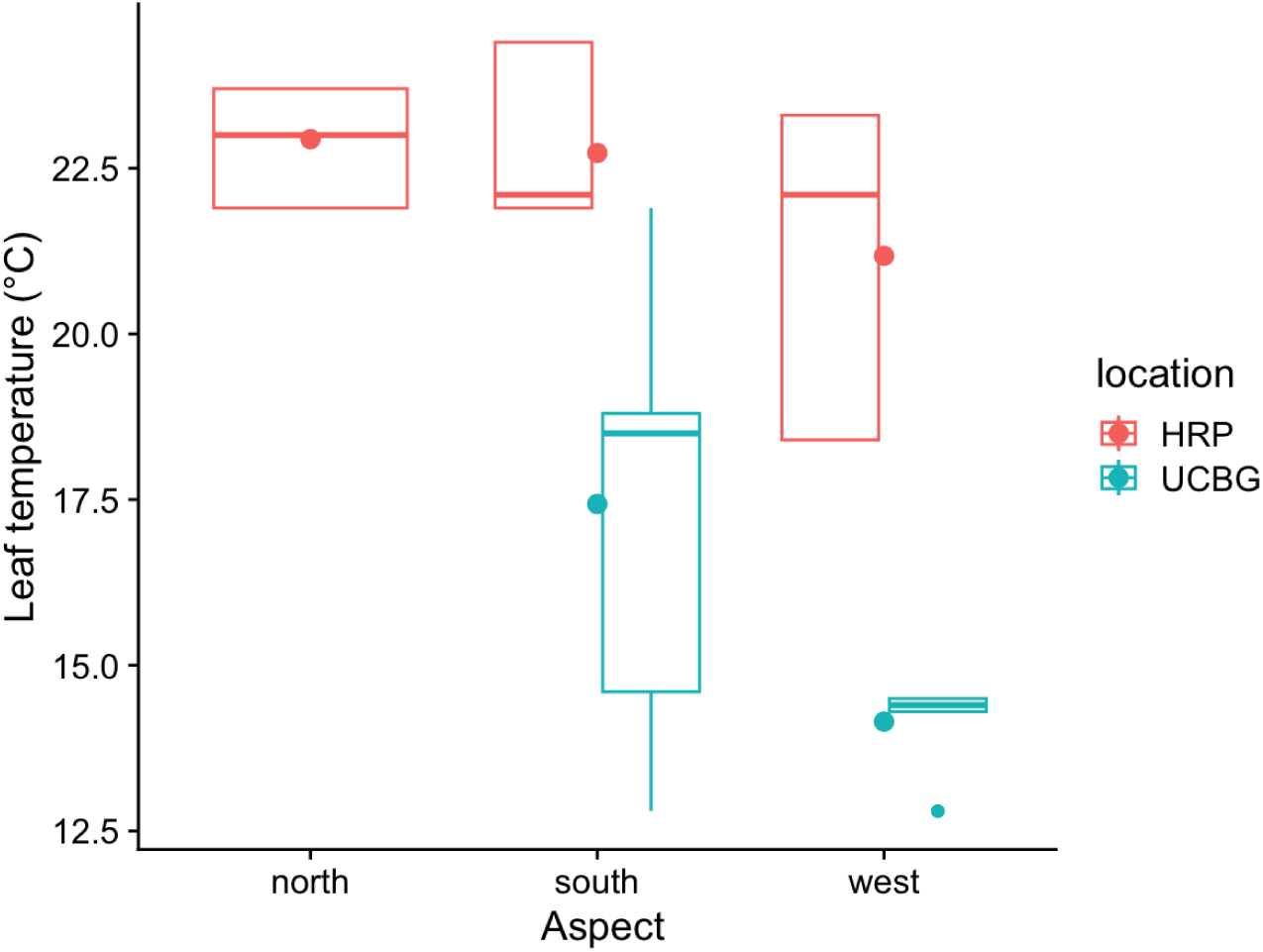
Leaf temperature across aspects at both sites.

**Figure 8.**
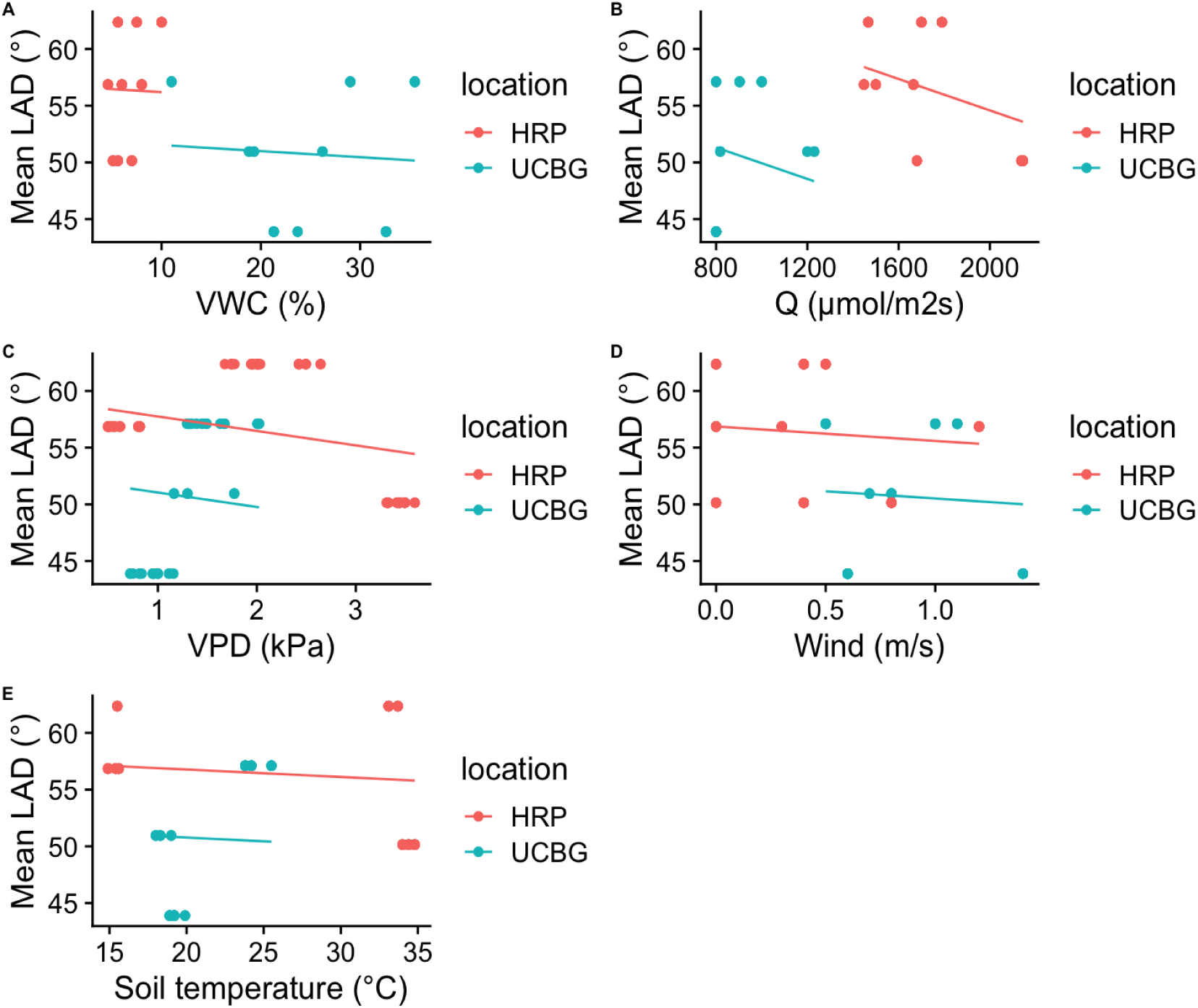
Microclimatic variables and mean LAD across sites. LAD means were statistically significantly different between sites. Mean values and standard deviations are found in Table 5 and Table 6.

**Figure 9.**
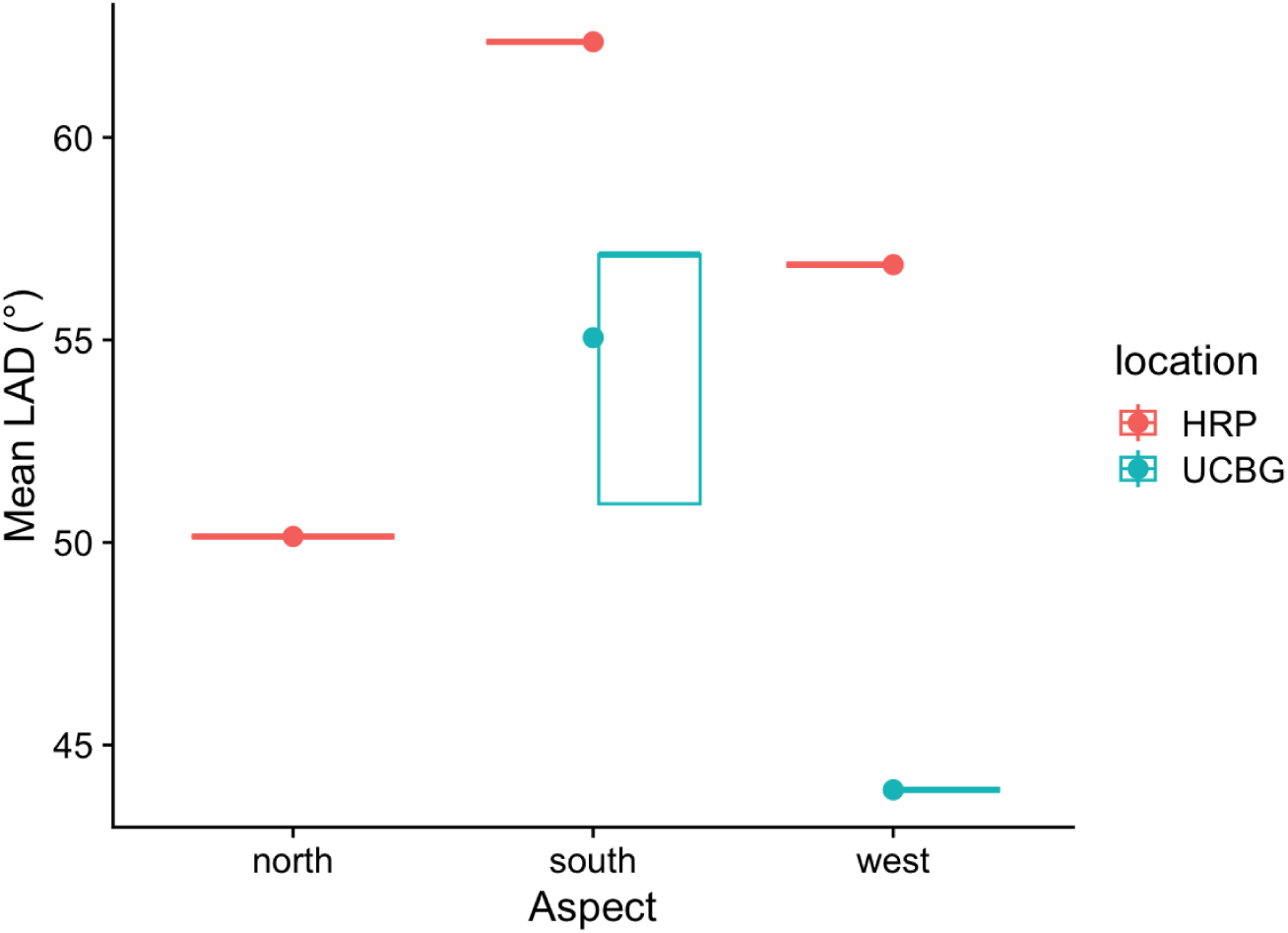
Mean LAD across aspects at both sites. LAD was significantly different between sites. At both sites, LAD was significantly different across aspects.

In line with my hypotheses, water availability and atmospheric demand, through VWC and VPD respectively, had some influence on plant functional traits. However, while the low VWC of HRP had little effect on photosynthesis (Figure 10, Table 9), there was a greater range of VPD conditions than at UCBG, with lower VPD earlier in the morning and greater VPD in the afternoon. Plants at HRP may obtain water from the saturated air early in the morning, when there is less sun and temperatures are cooler, and store that water instead of obtaining water from the low water-holding capacity soil.

**Figure 10.**
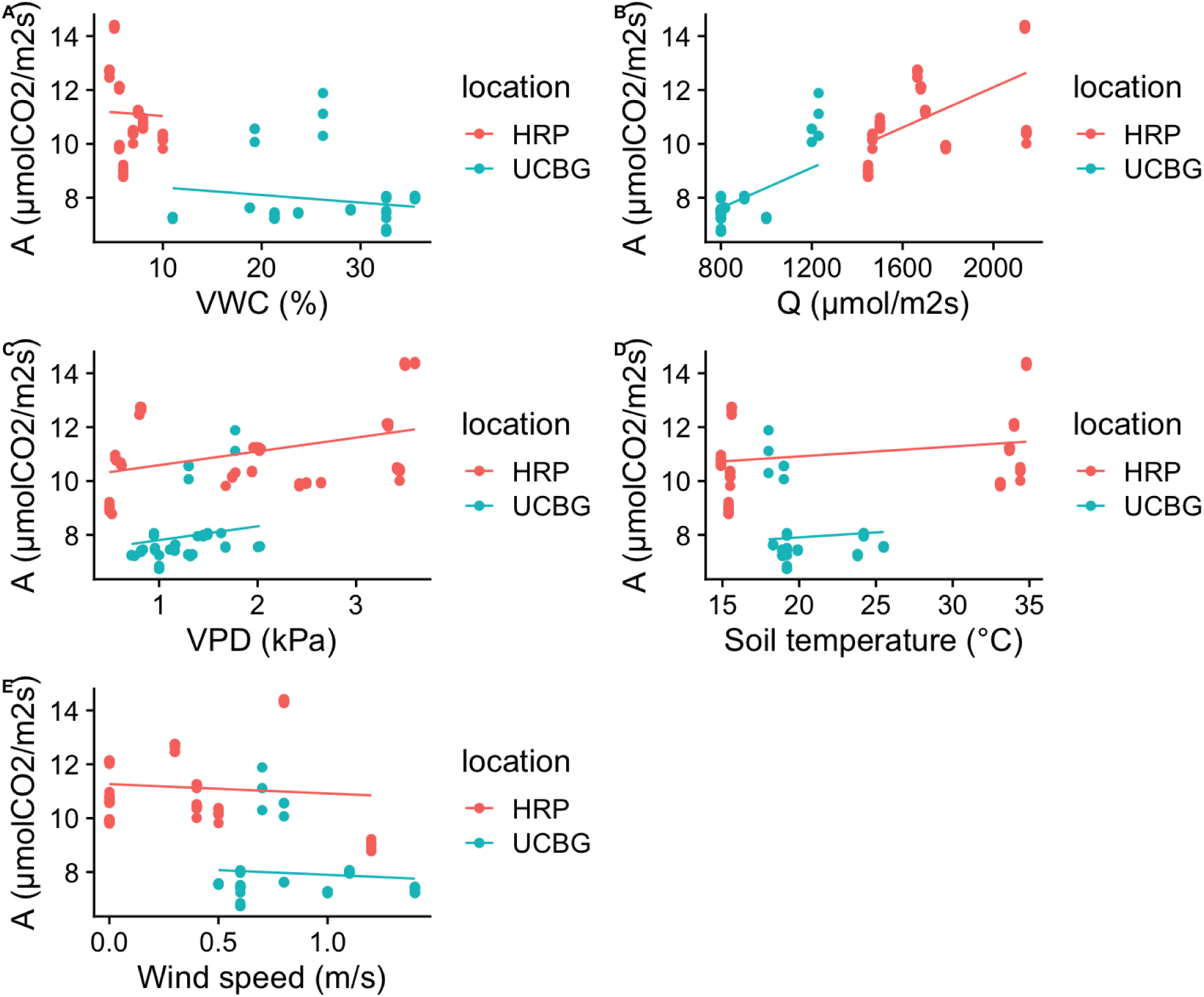
Microclimatic variables and photosynthesis across sites. Photosynthesis (A) means were statistically significantly different between sites. Mean values and standard deviations are found in Table 8.

**Figure 11.**
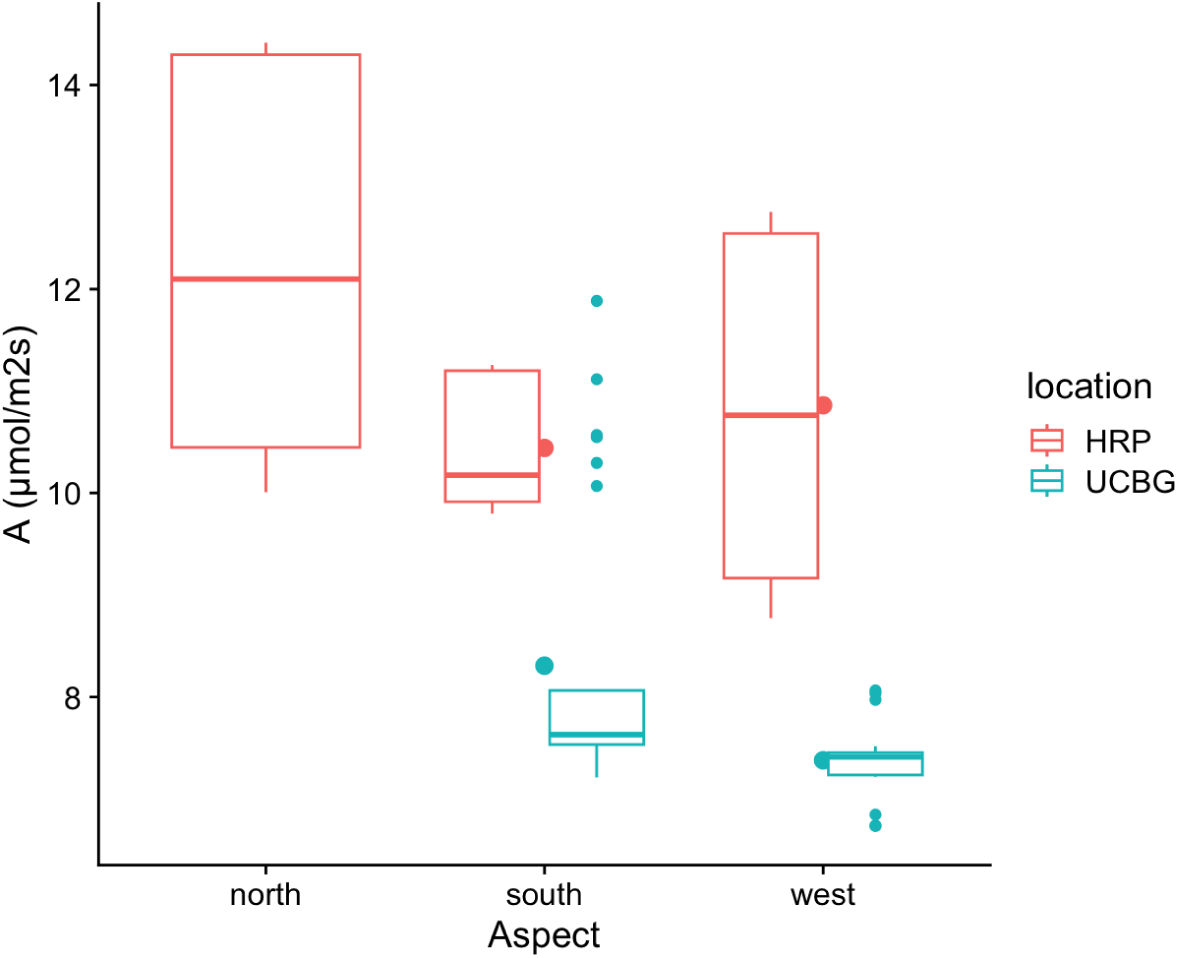
A across aspects at both sites. Photosynthesis (A) was significantly different between sites. At HRP, A differed significantly between north- and south- and between north- and west-facing slopes, but non significantly between south- and west-facing slopes. At UCBG, A was significantly different between aspects.

Plant productivity among sites reflected their environmental conditions. The strong, positive correlation between Q and A across many LMMs suggests that Q has the most influence on photosynthesis. At UCBG, there was lower solar radiation, increased VWC, and lower VPD; yet the mean carbon assimilation was lower than at HRP. This is somewhat contrary to my hypothesis that the site of higher solar radiation and more water availability will have greater photosynthesis, suggesting that solar radiation is a more influential predictor than water availability on the plants. *Arctostaphylos* are well-adapted to low soil moisture and high leaf temperature (Conard and Radosevich 1981) and are long considered drought-tolerant species, able to withstand high VPD during hot summers (Bhaskar and Ackerly 2006). Additionally, due to a greater range of VPD conditions and lower VWC, individuals at HRP may have acclimated or adapted to be more water use efficient (*W*i = *A*/*g*s) than individuals at UCBG (Lawson and Vialet-Chabrand 2019).

### Intraspecific Trait Variation

ITV is an important signifier in predicting plant response to changing climatic conditions as it captures the possible range of phenotypic responses in a species. In line with my hypotheses, my results showed that leaf morphological and plant physiological traits varied in response to microclimatic factors. Solar radiation and VPD significantly influenced leaf length, while VPD and VWC had strong predictor effects on LMA. Solar radiation, leaf temperature, and LAD had a strong influence on photosynthesis. The range of wind speed and soil temperatures at both sites had little effect on plant traits and ITV.

However, while these data also indicate strong site effects, there is little variation within sites, possibly due to a limited sample size and sampling season measured within each site. To capture a more complete picture of ITV within each site, it would be advantageous to increase sample sizes as well as measurement across seasons. If there is truly limited microclimate and ITV variation within each site, that could have strong effects on results, constraining the degree to which *A*. *crustacea* ssp. *crustacea* responds to abiotic and biotic conditions.

### Limitations and future directions

The timing of data collection presented potential biases in the data, with earlier collection at UCBG and later collection at HRP. This may have impacted the more temporary microclimatic conditions, like solar radiation, VPD, and wind speed. Alternatively, sampling these data over an extended time period may provide greater insight into trait and microclimate variation than data collected at a single time point in the growing season. Additional data collected across seasons would help clarify this point and would be valuable to do in the future.

Other factors affecting intraspecific trait variation within plants include genotype, ecosystem biotic dynamics, such as species competition and herbivory, and non-climatic abiotic factors, like nutrient availability (Roop et al. 2025). Due to limited time and resources, I did not consider these factors in my analysis. Future research can be done to include these factors in plant trait variation analysis. Variation in photosynthesis could also be a reflection of leaf age, local nutrient availability, or different physiological strategies intra-specifically (Longstreth and Nobel 1980, Wilson et al. 2001).

### Broader implications

Understanding plant functional traits provides important information to predict how plants will react to climate change. Assessing microclimate provides further, specific insight on the strength of certain microclimatic factors on *A*. *crustacea* ssp. *crustacea* traits. For example, solar radiation was the strongest predicting factor of all microclimatic conditions, while VPD may also be significant for water uptake. This information can be used to weigh these microclimatic factors more heavily than macroclimate when predicting plant response to more extreme climatic conditions. As temperatures, droughts, and wildfires in chaparral ecosystems increase, *Arctostaphylos* might respond in unpredictable ways and display increasing variability in their morphological and physiological traits. With higher ITV of a species, especially obligate seeder species in *Arctostaphylos*, there may be a greater likelihood of displaying adaptations to rapidly changing climates due to their genetic diversification (Kauffmann et al. 2021). Alternatively, if ITV is low, then these climatic stressors may further threaten brittleleaf manzanita and other endemic species, especially if microclimate conditions extend into new extremes. Studying the trait variation of a facultative seeder like *A*. *crustacea* ssp. *crustacea,* that typically clonally resprouts in the absence of fire, allows us to understand their functional response to current climate conditions and may aid in predicting resilience to future conditions.

However, in contrast to clonal propagation, genetic diversity within a population could potentially increase by encouraging and facilitating plant sexual reproduction with re-introduced managed fire regimes to brittleleaf and other manzanita habitats. A significant reduction in the regular intervals of controlled and wildfire in California habitats since colonization has been well documented, and only recently has western science begun to acknowledge the importance of fire in maintaining healthy chaparral habitats (Keeley et al. 2005, Underwood et al. 2022). As fire adapted plants, *Arctostaphylos* populations may benefit from a re-introduction of traditional and regular intervals of fire burning. Greater genetic diversity could therefore potentially cause a greater degree of ITV, thus potentially buffering plants from extreme climatic conditions.

Site-specific analysis of microclimate and plant health relationships also allows for better informed management strategies. Examining ITV across sites of different environmental conditions can better inform what the most optimal conditions for plant health are. Plant breeding programs targeted at plant restoration may consider examining microclimate, ITV, and genetic variation of manzanita post-fire to select for populations that may be better suited to future climate scenarios (Weiss et al. 2020). However, with further research done on the interaction of microclimate and ITV in manzanita species, it may be that ITV is already high enough to withstand future climates and no breeding program may be necessary.

## ACKNOWLEDGEMENTS

An enormous amount of gratitude, love, and appreciation to Dr. Roxy Cruz-de Hoyos, my mentor, role model, and the best plant ecophysiologist out there. Without her unwavering support, fierce determination, and encouragement, this thesis would not have been possible.

Equally large gratitude to Dr. Benjamin Blonder & the past and present members of the Macrosystems Ecology Lab for granting me access to the instruments used in this project, for offering helpful feedback, for teaching me so much about science, and for believing in me. My work with MEL has been supported by the USDA grant NIFA 2022-67019-36366.

Thank you to the UC Botanical Gardens, Clare Loughran, the East Bay Regional Park District (Permit 24-1252) and Michele Hammond for their support and allowing me to conduct research on their beautiful plants.

More gratitude and appreciation to Dr. Patina Mendez for her enthusiastic motivation and to the ES class of 2025 for their feedback, support, and collective suffering.

Finally, a huge thanks to my friends for always cheering me on and driving me and the big suitcases! Thank you to my lovely partner in crime for listening to my rants. Thank you to my mom for her support, my dad for teaching me to question, my siblings, and to my grandpa, the original plant person in our family. Special shout-out to Speckles!

